# Mechanistic Interpretability for Protein Language Models: A Validation Framework

**DOI:** 10.64898/2026.05.29.727021

**Authors:** Paul Chon, William B. Andreopoulos

## Abstract

Protein language models (PLMs) are shown to be powerful predictors of protein structure and function but their internal mechanisms remain poorly understood. Recent mechanistic interpretability methods have decomposed PLM representations into interpretable features, but they have not combined methods on a single biologically meaningful task. This paper tests whether an InterPLM sparse autoencoder and ProtoMech cross-layer transcoder can discover features in ESM-2 (6 layers, 8M) that can mainly discriminate between Class A *β*-lactamase and Class B *β*-lactamase with class C and D used as more challenging comparisons. The main goal is to find distinct features for Class A *β*-lactamase that are not shared by other classes. We find that both methods find distinct features for Class A *β*-lactamase, but the cross-layer transcoders show that the concepts for Class A *β*-lactamase seems to be distributed among nodes such as in layer 4 and 6 rather than one node. We also showcase a validation framework to prevent overclaiming the role of a node, and we use it to show that several strong nodes fail in some stages of the framework meaning that they cannot be the sole node that defines Class A *β*-lactamase.

## 1 Introduction

### 1.1 The Challenge of Antimicrobial Resistance

Antibiotic usage has become widespread in recent decades but antibiotics have become less effective. This is becoming an increasing health and economic concern with drug-resistant infections causing a significant increase in mortality and health care costs [1]. For example, estimates show that by 2050, over 10 million lives a year and a total of 100 trillion USD are at risk from drug-resistant infections [2].

This drug resistance, also called AMR (antimicrobial resistance), comes from a wide range of mechanisms but among these, hydrolysis (using water to break chemical bonds) of *β*-lactams by *β*-lactamases is the most common resistance mechanism for *β*-lactam, the most widely used antibiotic accounting for about 65 % of all antibiotic use [3], [4].

*β*-lactamases are enzymes that hydrolyze the *β*-lactam ring, which is the four-membered structure common in all *β*-lactam antibiotics [5]. By breaking this ring, *β*-lactamases render the antibiotic unable to bind its intended target, the penicillin-binding proteins (PBPs) responsible for bacterial cell wall cross-linking. The result is that bacteria carrying *β*-lactamase genes survive antibiotic exposure.

The Ambler classification system organizes *β*-lactamases into four molecular classes (A, B, C, and D) based on amino acid sequence homology (shared ancestry) [6]. Classes A, C, and D are serine *β*-lactamases: they use a serine in their active site to hydrolyze the *β*-lactam ring [7]. Class B enzymes are metallo-*β*-lactamases (MBLs): they require one or two zinc ions in their active site to hydrolyze the *β*-lactam ring [8].

To counteract the spread of AMR, detection systems or models can help in identifying novel variants for *β*-lactamases. Traditionally, tools such as BLAST and HMMER rely on pre-defined sequence patterns [9], [10]. However, this means that a novel enzyme could evade detection by having a completely different sequence that does not look like a typical enzyme. So, protein language models (PLMs) have been developed to take a different approach.

### 1.2 Protein Language Models in Biology

Protein language models (PLMs) learn embeddings, which are numbers that represent protein properties in a compact form, of protein sequences by training on large, unlabeled protein sequence databases [11]. Like natural language models trained on text, PLMs are trained to predict masked amino acids from their sequence context which makes the model learn the underlying syntax of protein sequences [12]. Through this training, the model develops an understanding of protein biology to cluster proteins with similar function or homology together without being given any pre-defined patterns [11], [13].

### 1.3 The Interpretability Problem

While PLMs achieve high accuracy on benchmark tasks such as structure prediction, their internal mechanisms remain hidden because they can contain millions or billions of parameters that would be difficult to keep track of and the embeddings that the PLMs output are not easily understood by humans. This complexity is why neural networks, including protein language models, are often described as black boxes [14]. Understanding what a PLM has learned and how it processes that information requires interpretability tools.

The field of mechanistic interpretability is trying to solve this gap in knowledge by developing methods to reverse engineer neural networks [15], [16]. For example, there are two main mechanistic interpretability methods that can open the black box of neural networks: sparse autoencoders (SAEs) analyze a single layer of a model [15] and cross-layer transcoders (CLTs) can look at multiple layers of a model [17]. Also, sparse autoencoders and cross-layer transcoders uncover interpretable features in neural networks which are essentially concepts or patterns that the model has learned [16]. Furthermore, cross-layer transcoders can compute circuits that represent a specified model behavior or concept through nodes that represent features discovered by cross-layer transcoders [17].

The gap in the current literature is that there is no work that combines both sparse autoencoders and cross-layer transcoders on a specific biological task.

The specific task in this paper is the discrimination of Class A *β*-lactamases from Class B *β*-lactamases with Class C and Class D used as more challenging comparisons [6]. This task is chosen because the goal is to find features for Class A *β*-lactamases that are not shared by Class B *β*-lactamases, which might be able to uncover new knowledge about Class A *β*-lactamases such as mechanisms or patterns. Also, the question becomes whether the discriminative features correspond to known functional difference or to a superficial pattern.

### 1.4 Contributions

The core claim of this paper is that we built and used a six-stage validation framework to clarify the true role of a node or feature, distinguishing a strong candidate node from a weak, superficial node. The main contributions of this paper are:

1. We apply two complementary mechanistic interpretability methods (InterPLM sparse autoencoders and ProtoMech cross-layer transcoders) to the same task, with both methods showing that there is Class A versus Class B discrimination in the PLM, ESM-2.
2. We evaluate the 16 node ProtoMech Class A circuit showing that the strongest nodes are in the early-layers (3 and 4) while there is only one strong candidate in the last layer.
3. We create and use a six stage validation framework (statistical benchmarking, cross-class comparison, family breadth, structural grounding, intervention controls, and metagenome comparison) on the Class A circuit.
4. We show the value of this framework through three main outcomes: cross-class comparison failure, superficial patterns, and mixed stage-dependent verdicts.

Figure 1 previews the six core assessment stages and the kinds of issues each stage can flag.

**Figure 1:**
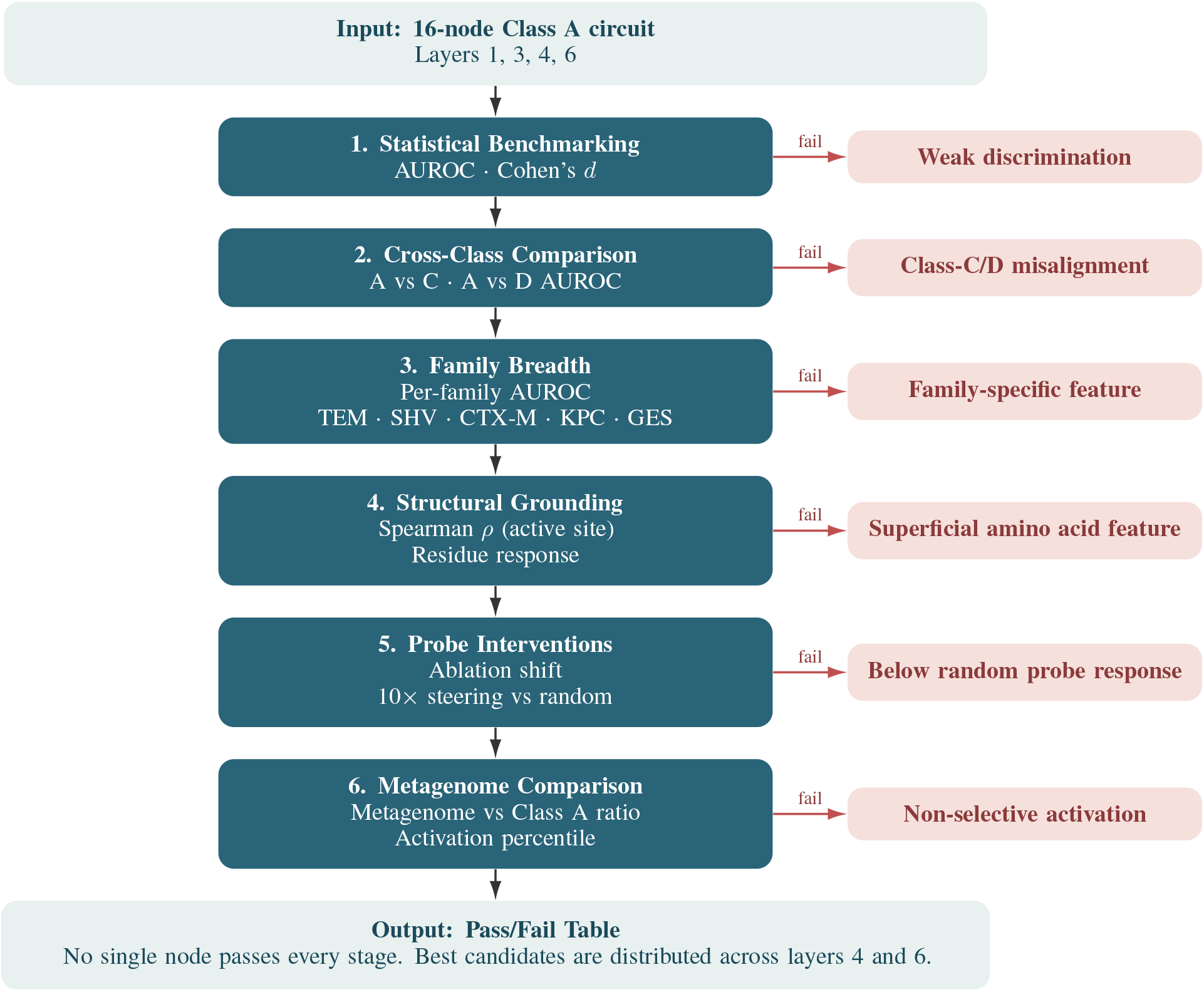
Sixteen candidate nodes from the ProtoMech Class A circuit enter at the top. Each stage applies a different assessment to the nodes and the red arrows pointing right are exits showing the kind of failures that each assessment can flag. Also, a node that fails one stage is still assessed at the other stages. Table 9 shows the pass or fail table summarizing the results of the nodes at each stage.

## 2 Background and Related Work

### 2.1 Beta-Lactamase Biology

Class A *β*-lactamases share several important recurring patterns of residues (amino acids incorporated into proteins) called motifs that provide a concrete reference to validate features from SAEs and CLTs [5], [3]:

- **SXXK motif** (positions 70-73), which contains a serine and a lysine involved in hydrolysis [7].
- **SDN motif** (positions 130-132), which helps stabilize the active site of the enzyme [7].
- **Omega loop** (positions 163-179), which is a motif that contains the residue Glutamate at position 166 that is found to be involved in hydrolysis [7].

### 2.2 Protein Language Models: ESM-2

This paper studies ESM-2, which is a family of protein language models, developed by Meta [18]. Specifically, this paper will look at the smallest version of ESM-2 with 8 million parameters and 6 layers called ESM-2 8M. ESM-2 8M takes in a protein sequence up to 1022 residues and outputs an embedding where each residue in the sequence has a vector of 320 dimensions. Also, each vector in this embedding will describe each residue’s properties using numbers that PLMs and even other regular machine learning models can understand. However, ESM-2’s internal mechanisms are hidden so we will use two main mechanistic interpretability tools to understand ESM-2: sparse autoencoders and cross-layer transcoders.

### 2.3 Sparse Autoencoders

Sparse autoencoders (SAEs) are motivated by the superposition hypothesis: neural networks often encode many features or concepts in a single neuron of a neural network [15]. SAEs address this by learning more features than there are neurons to reconstruct the original activation so that only a small number of features fire for any one input as shown in Figure 2 [15]. In effect, the method attempts to replace a dense representation with sparser, interpretable features. InterPLM applies this idea to ESM-2, producing SAEs for each layer of ESM-2 and found that ESM-2 contains interpretable concepts especially for biology related concepts [19]. Each InterPLM SAE contains 10,240 features which is a 32 × expansion of ESM-2’s hidden state of 320 dimensions [19]. For each input sequence, the SAE produces a vector of 10,240 dimensions at every sequence position [19].

**Figure 2:**
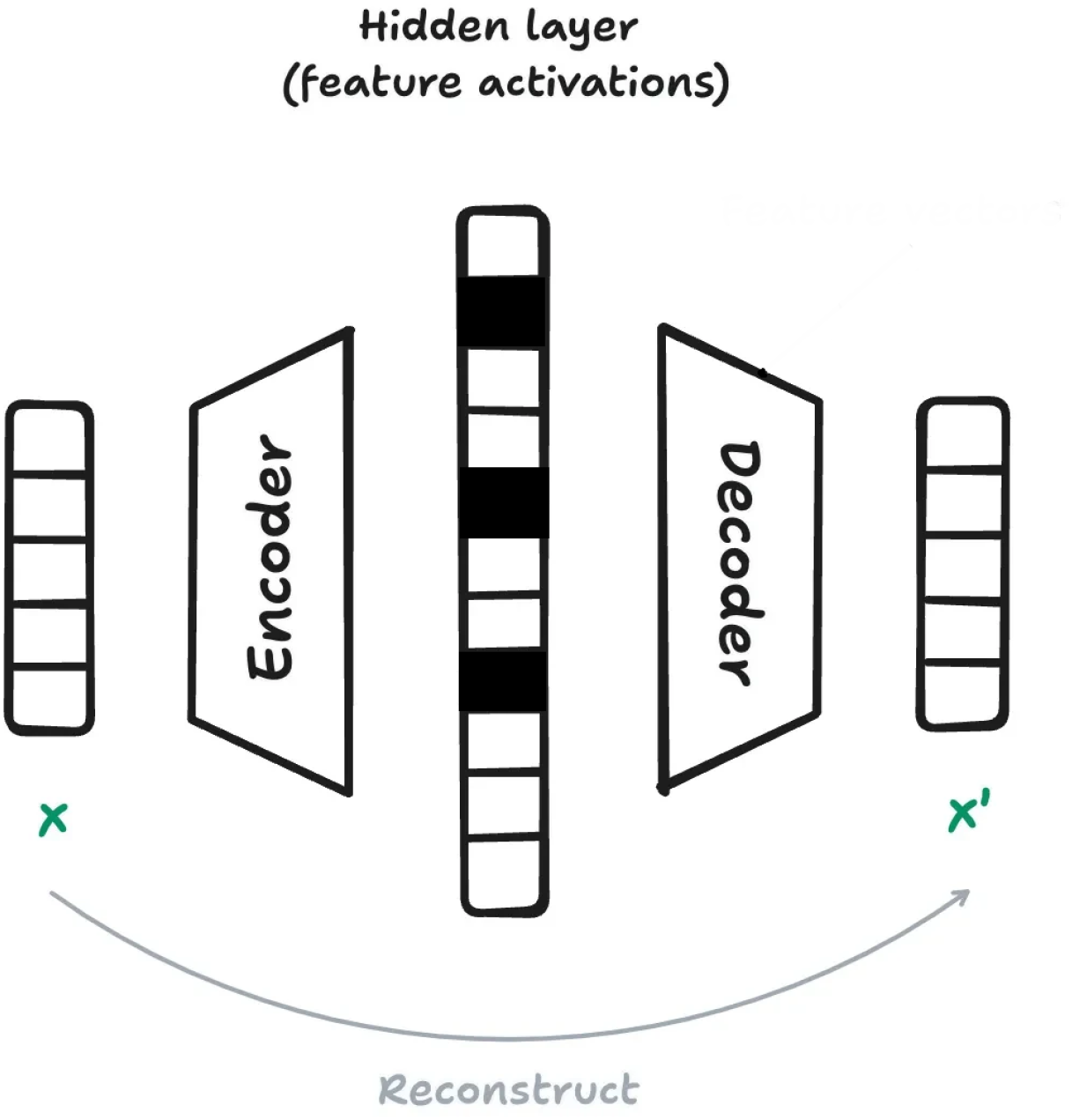
This figure shows how sparse autoencoders expand a layer into many features while encouraging only a few features to activate.

### 2.4 Cross-Layer Transcoders

On the other hand, cross-layer transcoders (CLTs) and the circuits computed from CLTs model how information is transformed across layers as shown in Figure 3 [17]. For example, ProtoMech trains a CLT on ESM-2 8M where the CLT is also six layers to match ESM-2 because the CLT acts as a replacement model for ESM-2. Specifically, the CLT replaces the multi-layer perceptron (MLP) component which is the same component as in large language models to process information and will output the embedding for ESM-2. Also, in this paper, nodes will sometimes be referenced with their layer number such as L5/1000 which stands for node 1000 in layer 5 of the CLT.

**Figure 3:**
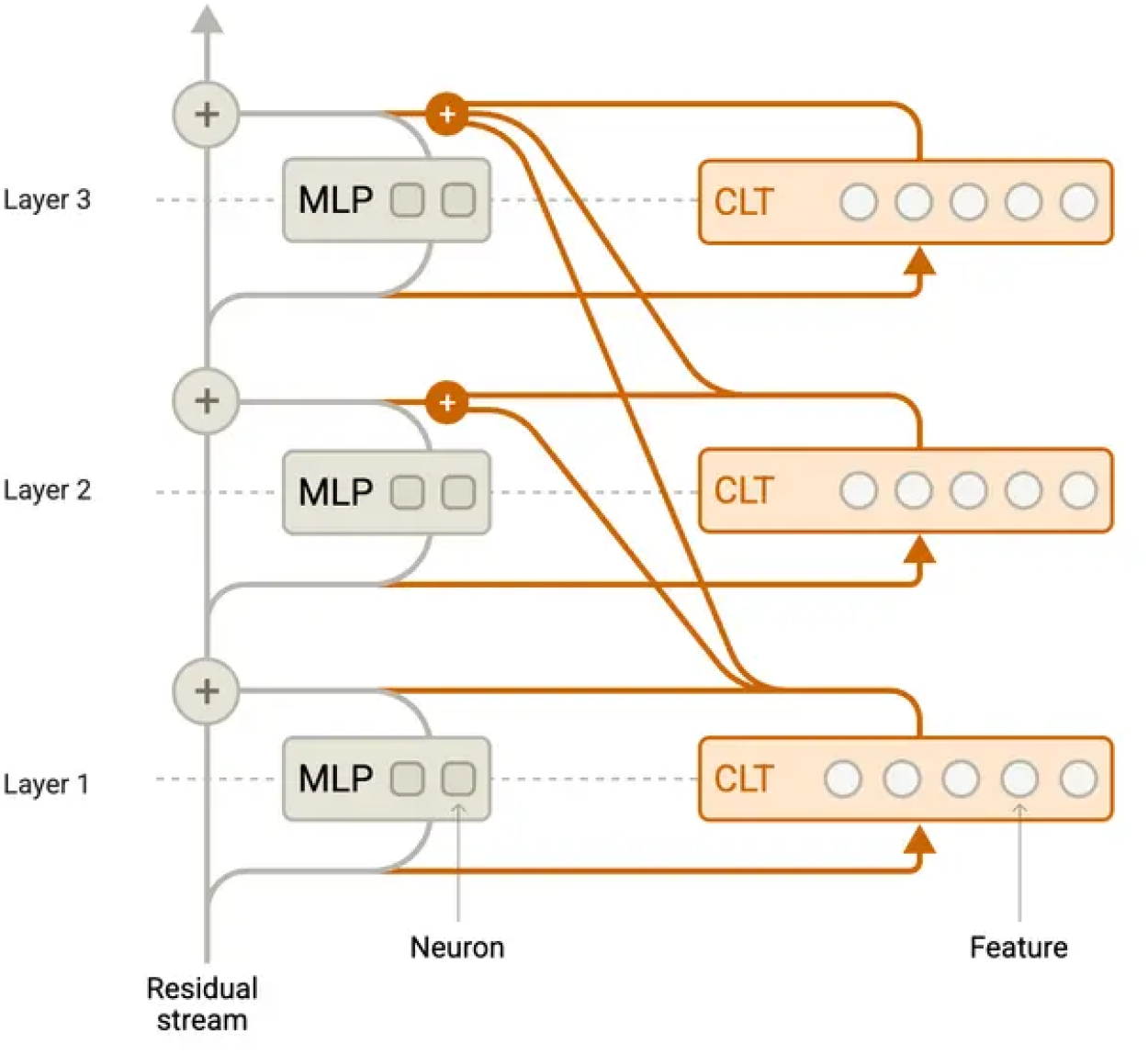
The CLT figure shows how a cross-layer transcoder maps information across layers and produces nodes or features in the replacement model [17]. It also shows how the CLT is replacing the MLP module of the model.

Also, ProtoMech computes circuits, which represent a specified model behavior or concept, by identifying the minimum amount of nodes needed to get to a similar level of performance as ESM-2 8M as shown in Figure 4 [17]. ProtoMech achieves up to 79% of model accuracy while using <1% of features discovered by the CLT [17].

**Figure 4:**
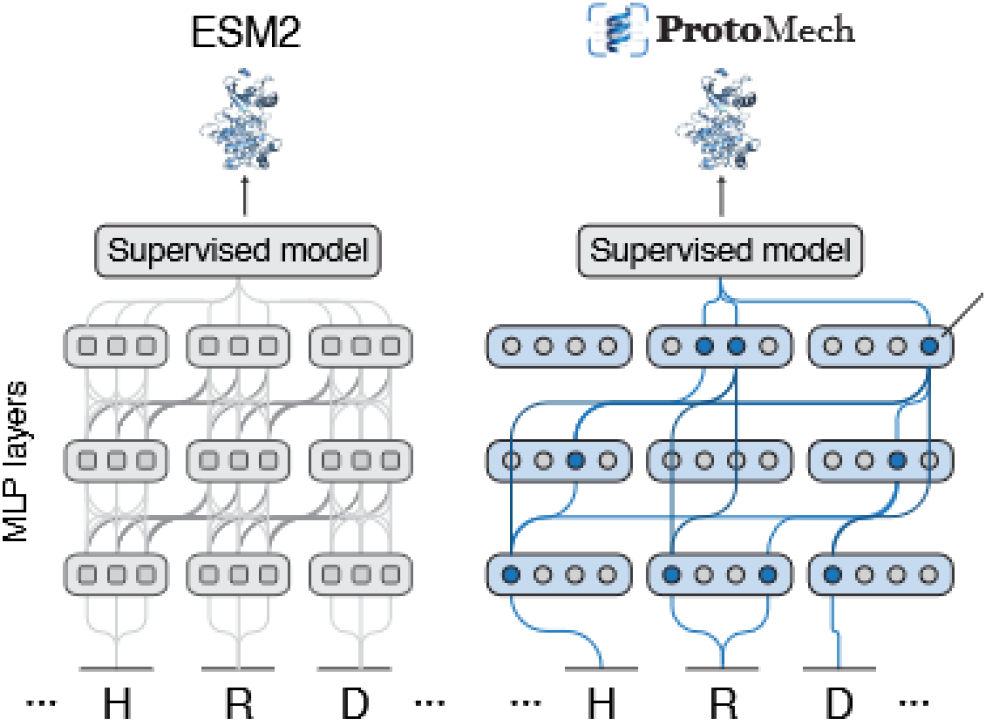
The figure shows how ProtoMech’s circuit discovery process identifies nodes that preserve the target model behavior [17].

### 2.5 Causal Intervention Methods

Beyond SAEs and CLTs, there is another set of mechanistic interpretability methods to validate their results: causal intervention which modifies a model’s internals to also understand how the model’s internals. This paper uses three causal intervention methods to validate that the features found are true: ablation, steering, and probe interventions.

Ablation lowers a feature’s activations to 0 and steering in this paper is only used to amplify a feature’s activations to see how the output of the model changes. For example, with a feature for Class A *β*-lactamase, ablating it should make the output farther away from Class A while steering and amplifying it should make the output closer to Class A. Both of these tools are used in probe intervention for CLTs, in which a linear probe or classifier is trained on the model’s activations and ablation and steering are applied onto target nodes or features.

## 3 Methods

### 3.1 Datasets

#### 3.1.1 Data Sources

*β*-lactamase sequences were collected from two primary databases: the Comprehensive Antibiotic Resistance Database (CARD) [20] and the InterPro database [21]. CARD was used as the main source of data while InterPro was used as a secondary source to add size to smaller *β*-lactamase classes such as Class B.

Sequences were assigned to Ambler classes [6] using name parsing, for example, TEM variants were assigned to Class A and NDM variants to Class B, CARD annotations, and InterPro family labels.

#### 3.1.2 Class A and B Family Distribution

The Class A dataset is dominated by eight families, with the five largest families (CTX-M, KPC, SHV, TEM, and GES) accounting for most of it. The most frequent Class B family labels were IMP, VIM, NDM, GOB, and CphA.

### 3.2 SAE Feature Extraction

#### 3.2.1 InterPLM SAE and Feature Analysis

For this paper, we used InterPLM’s layer 6 SAE trained on ESM-2 8M [19], [22]. To identify Class A and Class B SAE features, we computed: mean activation for Class A and Class B sequences, Welch’s *t*-test for the difference in the mean activations to give them class labels, and Cohen’s *d* to show how large the differences are. Features receive a class label, either Class A, Class B, or shared/other, using the sign of the mean difference together with the Welch test.

### 3.3 CLT Circuit Extraction

#### 3.3.1 ProtoMech CLT and Class A Circuit

We used ProtoMech’s cross-layer transcoder (CLT) [17] trained on ESM-2 8M [22]. The CLT contains 3200 nodes or features at each layer. Also, the main CLT analysis in this paper evaluates the 16 node ProtoMech Class A circuit. This circuit spans layers 1, 3, 4, and 6 and was recovered directly by ProtoMech’s circuit discovery on the CLT which seeks to use the minimum amount of nodes to recover original model behavior [17].

### 3.4 Validation Framework

This subsection details the six stages previewed in Figure 1.

The core metrics follow precedent from InterPLM and ProtoMech [17, 19]. Formal definitions of the metrics used throughout this subsection are given in Appendix subsection B.1.

#### 3.4.1 Circuit Benchmarking and Cross-Class Comparison

Each of the 16 Class A circuit nodes was benchmarked using per-sequence mean activation as the score for Class A versus Class B discrimination. The benchmark report includes AUROC, Cohen’s *d*, and cross-class comparison against Class C and Class D.

For the pass or fail column, a node passes benchmarking if its A vs B AUROC reaches 0.99 or higher, receives a mixed label between 0.75 and 0.99, and fails below 0.75. For cross-class comparison, a node passes if both A vs C and A vs D AUROC values are at least 0.90, receives a mixed label if one value falls between 0.75 and 0.90 while the other stays above 0.90, and fails otherwise.

#### 3.4.2 Family Breadth

In addition, to check whether a node relied too heavily on one dominant family type for Class A, we tested each of the five largest Class A families (TEM, SHV, CTX-M, KPC, and GES) individually against the Class B dataset. For each family, we measured how well the node could distinguish it from Class B. For family breadth, a node passes if its lowest per-family AUROC across the five families is 0.99 or higher, receives a mixed label if the lowest value is between 0.80 and 0.99, and fails below 0.80.

#### 3.4.3 Structural Grounding

Then, to assess whether nodes correspond to known Class A motifs or superficial patterns, we mapped high activating residues onto Class A structures predicted with ESMFold [22]. For each node, we recorded: (1) the Spearman correlation and p-values to find whether it concentrates on or near the enzyme’s active site [3], [7], and (2) whether it’s simply responding to residues rather than true structural features. For the Spearman correlation values, a negative *ρ* value means a node is concentrated near the enzyme’s active site while zero is weak and positive means the node is not concentrated near the active site. Also, the p-value helps to test whether the Spearman correlation value is real rather than happening by chance with a lower p-value meaning that there’s a low chance the value happened by chance.

For structural grounding, a node fails if a single amino acid or residue is in seven or more of the node’s ten highest activating positions which shows a feature for a single amino acid rather than a true structural feature. A node passes if its Spearman *ρ* is negative with *p <* 0.05, meaning its activations concentrate near the active site and the results did not happen from random chance. A node receives a mixed label if it avoids the superficial amino acid pattern but its active site distance is weak or not significant.

#### 3.4.4 Probe Interventions

Probe interventions were run separately for each circuit layer. For a target layer *L*, a logistic-regression probe was trained on that layer’s mean-pooled CLT representation to distinguish Class A from Class B. Each target node in that layer was then perturbed in two ways:

- **Zero ablation**: set the target node to zero before recomputing the probe response.
- **10**× **steering**: multiply the target node by 10 in the steered class and measure the induced shift.

For comparison, we tested random control nodes alongside each real node. We measured how much each intervention shifted the classification prediction compared to the random baseline which is represented by the ablation logit shift and the steering probability shift. These tests show which nodes matter most for discriminating Class A vs Class B. A node passes if both its ablation and 10 × steering z-scores are 1 standard deviation above the random controls, receives a mixed label if at least one z-score is positive but both z-scores do not go above 1, and fails if both z-scores are zero or negative.

#### 3.4.5 Metagenome Validation

The purpose of this stage is to test the nodes on a different, real-world dataset other than the one we used in the beginning to find the Class A circuit. We scanned all 16 Class A circuit nodes across 5000 proteins from the NMPFamsDB metagenome dataset [23]. For each node, we had two assessments: the first assessment measured how strongly it activates on the metagenome sequences compared to the Class A sequences and the second assessment looks at how many metagenome sequences exceed each node’s 75th, 90th, and 95th percentile threshold for activation (this is a specific activation value different for each node where 75%, 90%, or 95% of the node’s activations for the Class A sequences are at or below it). The first assessment explains whether the nodes are looking only at Class A sequences, meaning a low ratio of metagenome vs Class A sequences, or are looking at other superficial patterns, meaning a high ratio of metagenome vs Class A sequences. The second assessment shows whether a node might be a true Class A feature with a low amount of metagenome sequences above the thresholds or a superficial pattern with a high amount of metagenome sequences above the thresholds because the metagenome dataset does not contain all *β*-lactamases. A node passes if its metagenome vs Class A ratio is below 0.10 and no metagenome sequences exceed the node’s 95th percentile threshold. A node receives a mixed label if the ratio stays below 0.50 and at most 5 sequences exceed the 95th percentile, otherwise the node fails.

## 4 Results

### 4.1 SAE Features Distinguish Class A and Class B *β*-Lactamases

In this first step, we looked at the InterPLM SAE’s layer 6 activations from all Class A and Class B sequences. Of the 10,240 SAE features in layer 6, 5,890 showed a meaningful difference in mean activations from the Welch’s *t*-test: 3,288 features were labeled Class A and 2,602 were labeled Class B. Also, the strongest individual features showed large Cohen’s *d*. Table 4 reports the eight features with the largest absolute Cohen’s *d* values. Figure 5 visualizes the top eight layer 6 SAE features by their absolute Cohen’s *d* value together with the family composition of those features.

**Table 1:** Dataset Composition.

| Dataset | N | Source | Use |
| --- | --- | --- | --- |
| Class A (serine BLAs) | 1,152 | CARD + InterPro | Primary positive class |
| Class B (metallo BLAs) | 347 | CARD + InterPro | Primary negative class |
| Class C (AmpC) | 1,451 | CARD + InterPro | Cross-class comparison |
| Class D (OXA) | 1,272 | CARD + InterPro | Cross-class comparison |

**Table 2:** Class A Family Breakdown.

| Family | N | Clinical Significance |
| --- | --- | --- |
| CTX-M | 267 | Extended-spectrum; most prevalent globally |
| KPC | 229 | Carbapenemase; major clinical concern |
| SHV | 216 | Commonly plasmid-mediated |
| TEM | 216 | Most common in Enterobacteriaceae |
| GES | 64 | Variable-spectrum; some carbapenemase activity |
| CARB | 54 | Carbenicillin-hydrolyzing |
| VEB | 39 | Extended-spectrum |
| IMI | 28 | Carbapenemase |

**Table 3:** Class B Family Breakdown.

| Family | N | Clinical Significance |
| --- | --- | --- |
| IMP | 100 | Imipenem-hydrolyzing; often integron-associated |
| VIM | 90 | Verona integron-encoded MBL |
| NDM | 70 | New Delhi MBL; major clinical emergency |
| GOB | 53 | Primarily found in *Flavobacterium* spp. |
| CphA | 9 | Narrow-spectrum carbapenemase |

**Table 4:**
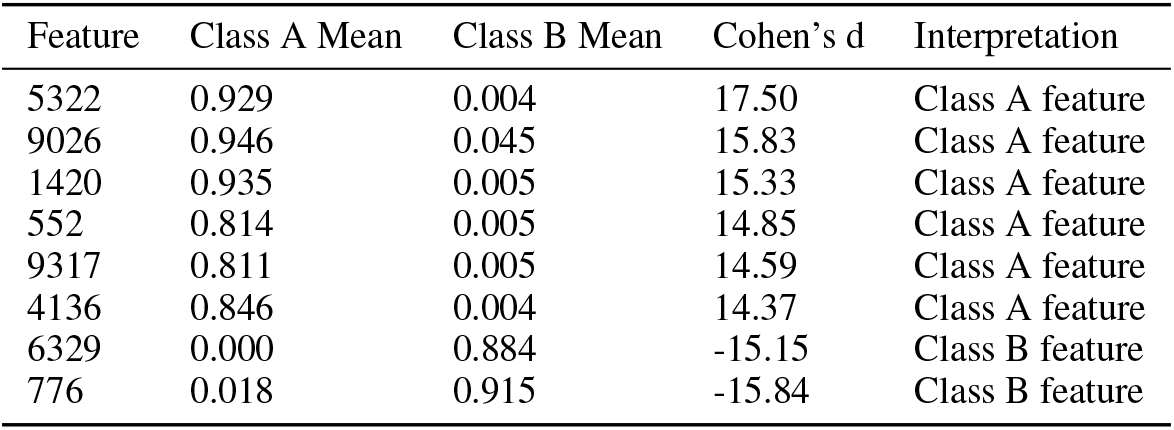
Top SAE features by Cohen’s *d* value.

**Figure 5:**
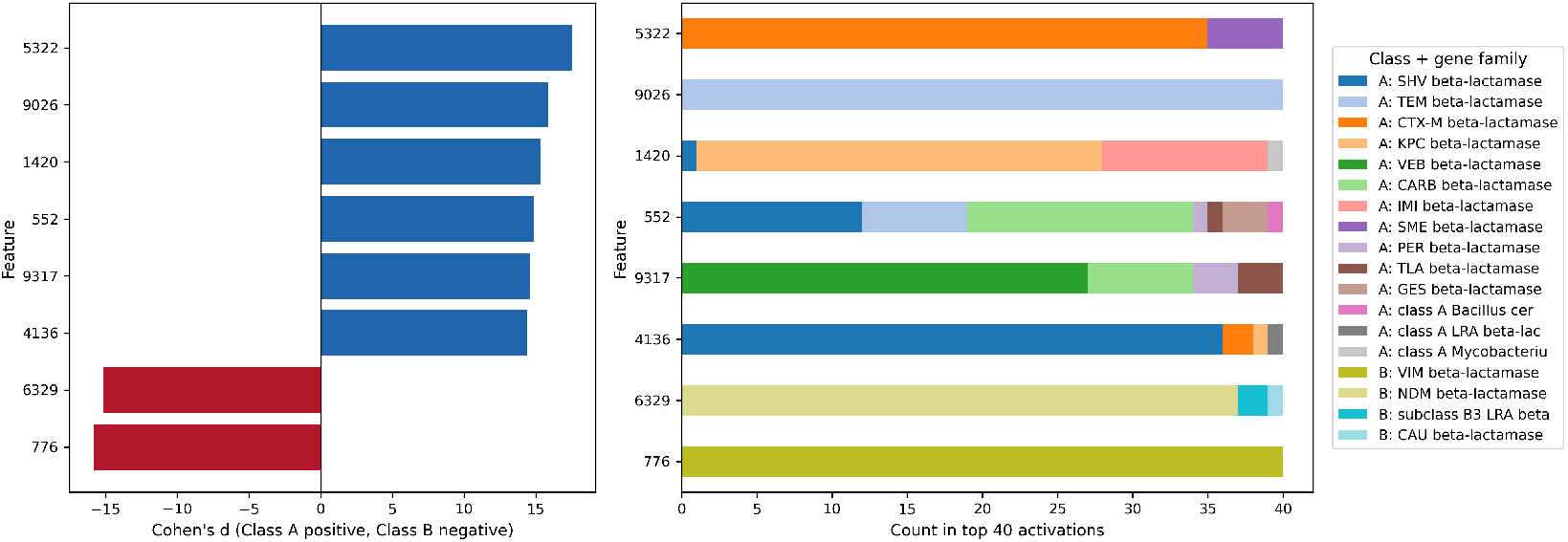
The left figure shows the Cohen’s *d* value for the top eight SAE features. The right figure shows the families for the top 40 sequences that have the highest activations for the top eight SAE features.

We see in the left figure of Figure 5, layer 6 SAE features show strong class separation in ESM-2, with large Cohen’s *d* for both Class A and Class B features. Also, in the right figure of Figure 5, most of these top features, except for Feature 776, also show that they represent several different families within their classes so they are not dependent on one family. These results using InterPLM’s layer 6 SAE show that ESM-2 does contain features that can cleanly separate Class A and Class B, so now we look to see if this result holds in CLTs as well, specifically the Class A circuit discovered by ProtoMech’s circuit discovery process [17].

### 4.2 Class A Circuit Benchmarking and Cross-Class Comparison

The Class A circuit contains 16 nodes spanning layers 1, 3, 4, and 6. The results of benchmarking the circuit ordered by rank, which is determined by how much a node attributes to the output, is shown in Table 5 and Figure 6. Appendix Table 11 provides the nodes ordered by layer. The results show that seven of the top eight circuit nodes lie in layers 3 or 4, while L6/1577 ranks eighth within the circuit and leads the layer 6 group. So, the strongest nodes are concentrated in these middle layers of the CLT. For example, L4/543 and L4/2576 both reach a A vs B AUROC (Area Under the Receiver Operating Characteristic Curve) of 1.0, which means that it is perfect classifier or discriminator, with strong cross-class results as well.

**Table 5:**
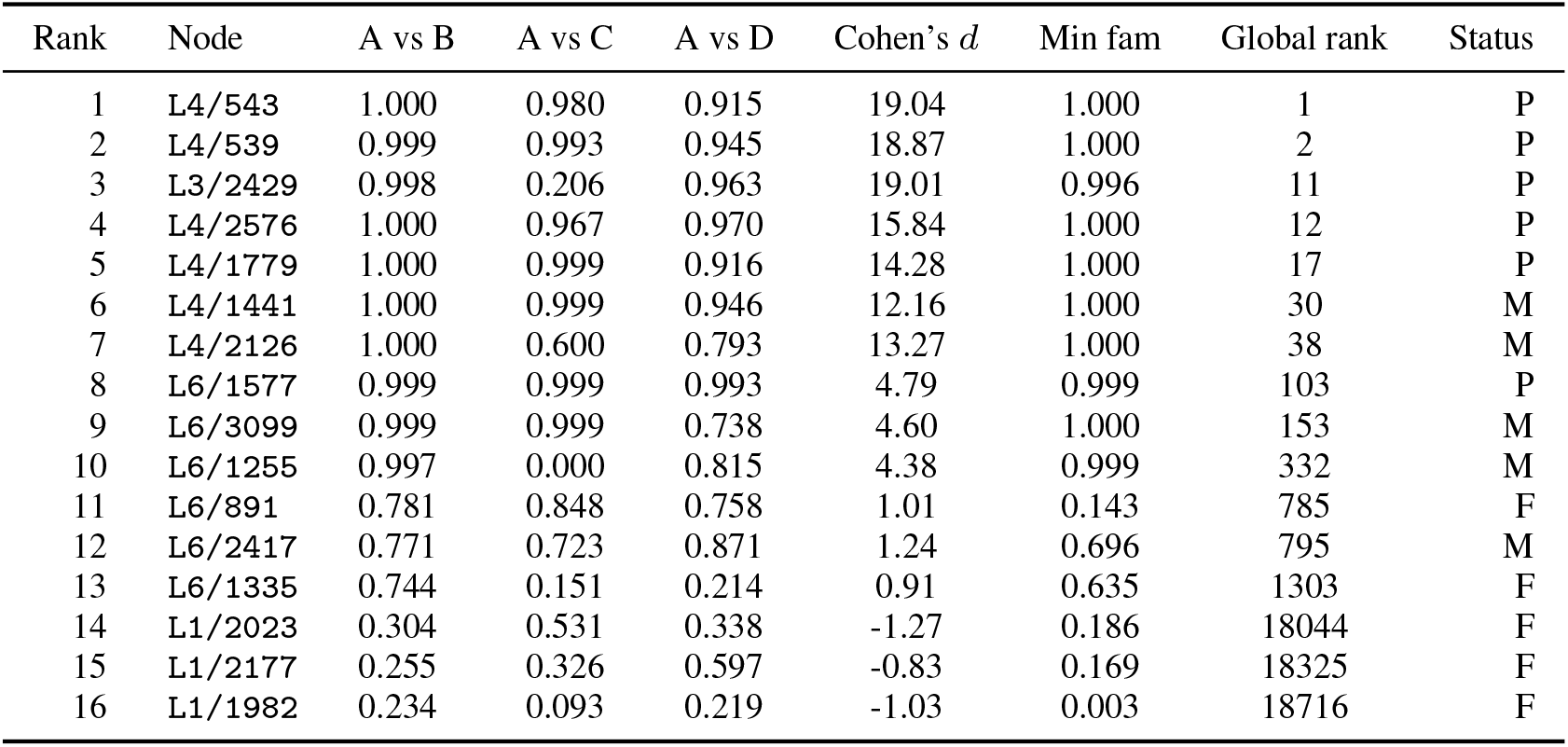
Full Class A circuit benchmark summary, ordered by rank.

| Rank | Node | A vs B | A vs C | A vs D | Cohen's $d$ | Min fam | Global rank | Status |
| --- | --- | --- | --- | --- | --- | --- | --- | --- |
| 1 | L4/543 | 1.000 | 0.980 | 0.915 | 19.04 | 1.000 | 1 | P |
| 2 | L4/539 | 0.999 | 0.993 | 0.945 | 18.87 | 1.000 | 2 | P |
| 3 | L3/2429 | 0.998 | 0.206 | 0.963 | 19.01 | 0.996 | 11 | P |
| 4 | L4/2576 | 1.000 | 0.967 | 0.970 | 15.84 | 1.000 | 12 | P |
| 5 | L4/1779 | 1.000 | 0.999 | 0.916 | 14.28 | 1.000 | 17 | P |
| 6 | L4/1441 | 1.000 | 0.999 | 0.946 | 12.16 | 1.000 | 30 | M |
| 7 | L4/2126 | 1.000 | 0.600 | 0.793 | 13.27 | 1.000 | 38 | M |
| 8 | L6/1577 | 0.999 | 0.999 | 0.993 | 4.79 | 0.999 | 103 | P |
| 9 | L6/3099 | 0.999 | 0.999 | 0.738 | 4.60 | 1.000 | 153 | M |
| 10 | L6/1255 | 0.997 | 0.000 | 0.815 | 4.38 | 0.999 | 332 | M |
| 11 | L6/891 | 0.781 | 0.848 | 0.758 | 1.01 | 0.143 | 785 | F |
| 12 | L6/2417 | 0.771 | 0.723 | 0.871 | 1.24 | 0.696 | 795 | M |
| 13 | L6/1335 | 0.744 | 0.151 | 0.214 | 0.91 | 0.635 | 1303 | F |
| 14 | L1/2023 | 0.304 | 0.531 | 0.338 | -1.27 | 0.186 | 18044 | F |
| 15 | L1/2177 | 0.255 | 0.326 | 0.597 | -0.83 | 0.169 | 18325 | F |
| 16 | L1/1982 | 0.234 | 0.093 | 0.219 | -1.03 | 0.003 | 18716 | F |

**Figure 6:**
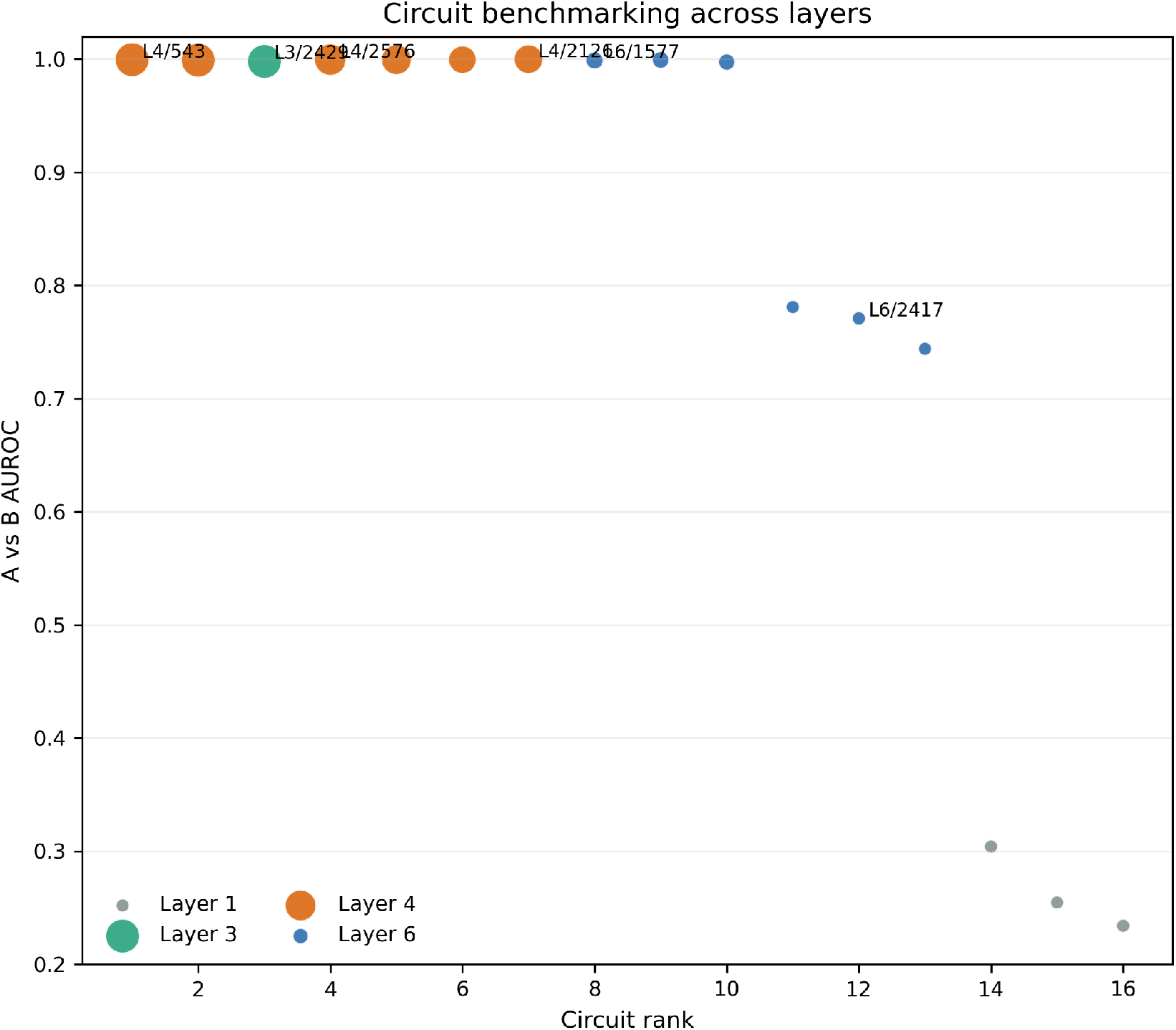
The figure shows A vs B AUROC across the 16-node Class A circuit, with point size scaled by Cohen’s *d* and color indicating layer. Also, the figure shows that the best nodes are in the middle layers specfically in layer 4 while the worst nodes are in layer 1 and some in layer 6.

On the other hand, the results show that a high A vs B performance is not enough to call a node a strong candidate. L3/2429 is ranked third because of its high AUROC value of 0.998 and large Cohen’s *d* value of 19.01, but has a A vs C AUROC of 0.206 which means that this node cannot separate between Class A and Class C. So, this stage of the validation framework is able to catch nodes that represent features that are not distinctly for Class A but might be features shared between the classes.

### 4.3 Family Breadth

The results for Class A family breadth is visualized in Figure 7 and the results are also in Appendix Table 12. The results show that the best nodes are not dependent on a single family. For instance, the top two nodes, L4/543 and L4/539, all achieve perfect or near-perfect AUROC values across all the Class A *β*-lactamase families. So, strong family breadth does not belong to one layer or one node.

**Figure 7:**
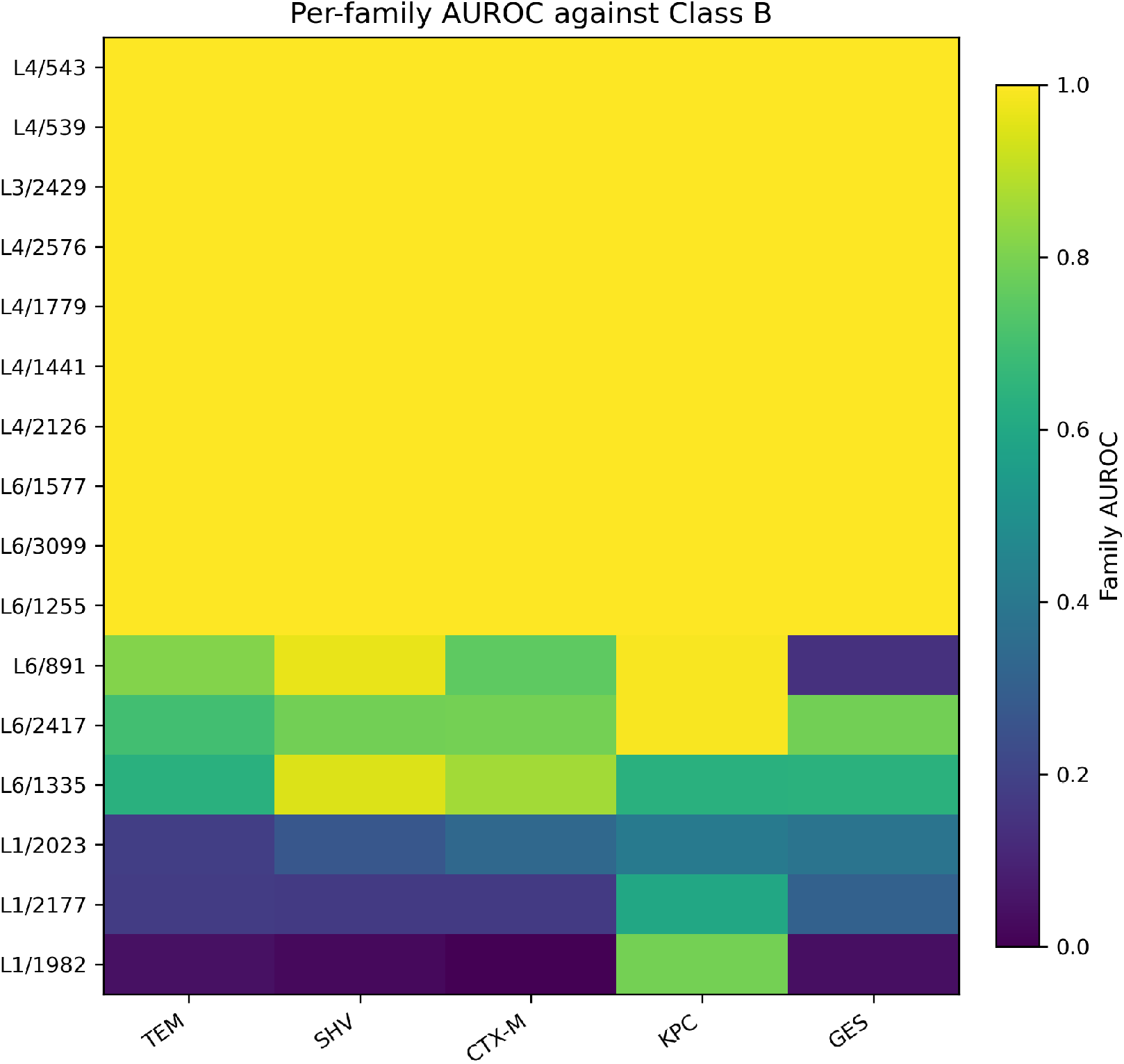
The figure shows per-family AUROC against Class B for the Class A circuit.

However, the results do show that the later layers tend to be better discriminators than the early layers because none of the nodes in layer 1 get an AUROC near 1.0 except for L1/1982 with the KPC family. On the other hand, even the weaker nodes in layer 6 achieve an AUROC of 0.5 or higher for all Class A families except for L6/891 on the GES family.

### 4.4 Structural Grounding

The structural grounding results are shown in Table 6. The strongest node in the circuit for this stage is L4/1779 with the highest Spearman correlation value of *ρ* = − 0.365 and one of the lowest p-values of 2.52 × 10^−08^. These values mean that L4/1779 is the most concentrated at or near the active site from the *ρ* value and that the Spearman correlation value is significant and not random chance from the p-value. L4/2576 is similarly strong with a *ρ* of -0.335 and the lowest p-value of 9.7 × 10^−10^, while the top ranked node for the Class A circuit L4/543 has weaker results of *ρ* = − 0.173 and *p* = 0.0047.

**Table 6:** Structural grounding summary for all Class A circuit nodes.

| Rank | Node | $\rho_{cat}$ | $p$ | Top res. | Count | Top-10 residues | Status |
| --- | --- | --- | --- | --- | --- | --- | --- |
| 1 | L4/543 | -0.173 | 0.00474 | L | 3 | WLELENRRLS | P |
| 2 | L4/539 | 0.014 | 0.826 | V | 2 | VPVARNDPGW | M |
| 3 | L3/2429 | -0.198 | 0.00119 | L | 5 | LLGLFQLYLY | P |
| 4 | L4/2576 | -0.335 | 2.52e-08 | V | 5 | VLAVVLVAPV | P |
| 5 | L4/1779 | -0.365 | 9.74e-10 | V | 3 | LVAVGTVLAA | P |
| 6 | L4/1441 | -0.019 | 0.755 | V | 4 | LAVVGPVIVY | M |
| 7 | L4/2126 | -0.068 | 0.268 | G | 2 | GKDWVARVYG | M |
| 8 | L6/1577 | -0.121 | 0.0488 | G | 2 | VGINGPHNRW | M |
| 9 | L6/3099 | 0.081 | 0.189 | L | 4 | LLILGLNTTG | M |
| 10 | L6/1255 | -0.047 | 0.445 | Q | 2 | QNEFLPQPRA | M |
| 11 | L6/891 | -0.082 | 0.182 | L | 10 | LLLLLLLLLL | F |
| 12 | L6/2417 | 0.051 | 0.41 | A | 10 | AAAAAAAAAA | F |
| 13 | L6/1335 | 0.049 | 0.424 | Q | 9 | QQQQQQQQQ | F |
| 14 | L1/2023 | -0.025 | 0.687 | V | 10 | VVVVVVVVVV | F |
| 15 | L1/2177 | 0.127 | 0.0398 | G | 10 | GGGGGGGGGG | F |
| 16 | L1/1982 | -0.002 | 0.969 | Q | 5 | QWQSNNQQQS | F |

Figure 8 visualizes which nodes might only be responding to residues. For the top 10 highest activating residues, L6/2417 only has alanine, L6/891 only has leucine, and several of the layer 1 nodes only have a single residue. So, these nodes fail in the structural grounding stage which like previous stages tends to happen more with layer 1 nodes.

**Figure 8:**
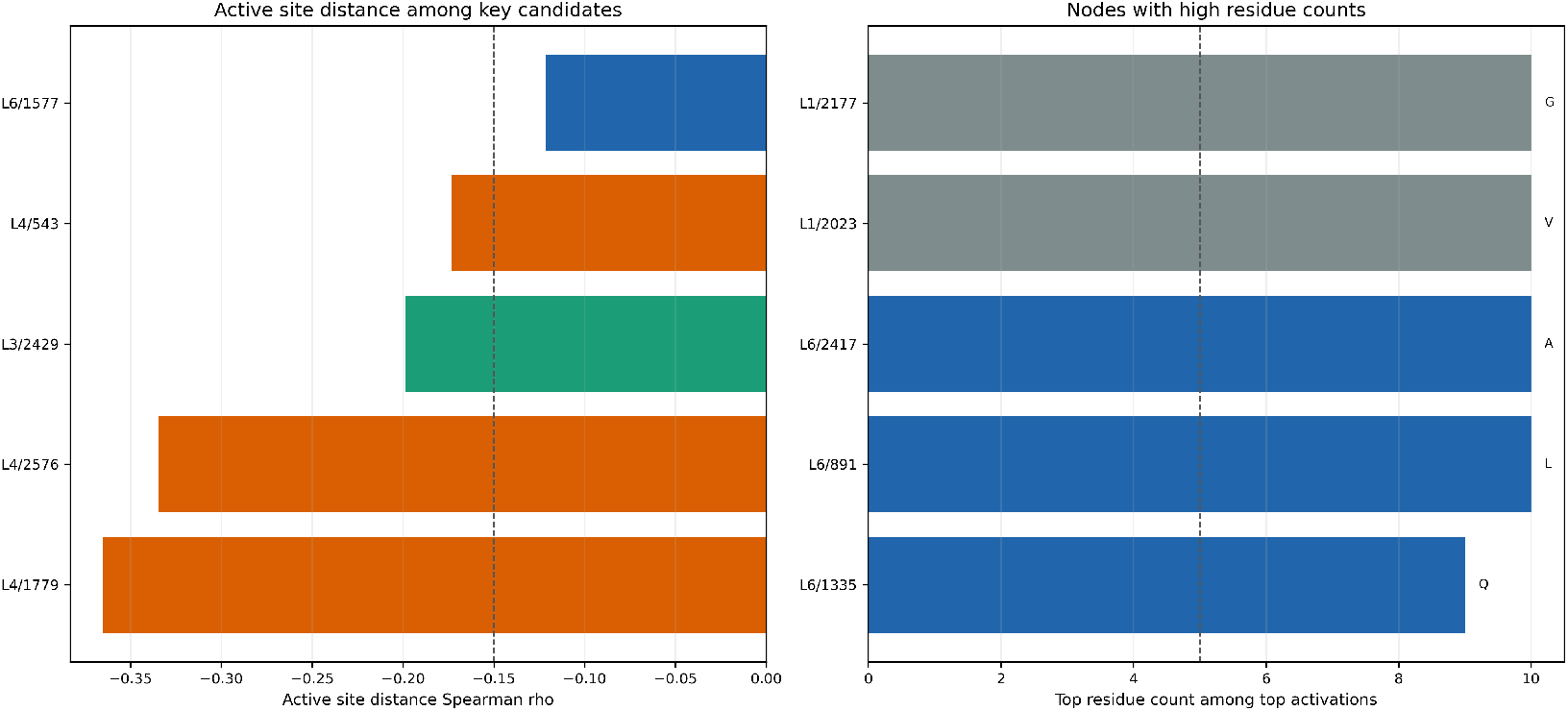
The left figure shows the Spearman’s correlation value *ρ* for the nodes that have high *ρ* and low p-value meaning they concentrate on the active site and are real values not found by random chance. The right figure shows the counts for the top residue for nodes that have high counts.

Also, in this stage, L3/2429 has good results for *ρ* and p-value, and doesn’t show any signs of only responding to residues which means it passes structural grounding. However, it failed on cross-class comparison so this shows that every stage of the validation framework matters to not jump to conclusions.

### 4.5 Probe Interventions

Table 7 shows the probe intervention results and their exact values. Figure 9 visualizes the probe intervention results measured by standard deviations above the random control results. The results show that all six layer 4 nodes output negative standard deviations on both intervention summaries. For example, L4/543 scores −0.52 standard deviations on ablation and −0.45 standard deviations on 10 × steering, while L4/2576 scores − 0.52 standard deviations and − 0.54 deviations respectively. These values show that the layer 4 nodes performed worse than the random control nodes, so the layer 4 nodes fail the intervention stage.

**Table 7:** Probe intervention summary for all Class A circuit nodes.

| Rank | Node | Ablation z | Steer z | $\vert \Delta \logit\vert $ | $\vert \Delta p\vert $ | Status |
| --- | --- | --- | --- | --- | --- | --- |
| 1 | L4/543 | -0.52 | -0.45 | 0.125 | 2.39e-05 | F |
| 2 | L4/539 | -0.50 | -0.49 | 0.127 | 2.13e-05 | F |
| 3 | L3/2429 | 0.52 | 1.79 | 0.178 | 0.000137 | M |
| 4 | L4/2576 | -0.52 | -0.54 | 0.124 | 1.71e-05 | F |
| 5 | L4/1779 | -0.47 | -0.49 | 0.117 | 1.8e-05 | F |
| 6 | L4/1441 | -0.47 | -0.51 | 0.116 | 1.62e-05 | F |
| 7 | L4/2126 | -0.53 | -0.52 | 0.124 | 1.87e-05 | F |
| 8 | L6/1577 | 1.16 | 3.24 | 0.389 | 8.62e-05 | P |
| 9 | L6/3099 | 1.01 | 1.57 | 0.315 | 0.000149 | P |
| 10 | L6/1255 | 3.70 | 8.50 | 0.338 | 9.13e-05 | P |
| 11 | L6/891 | -2.44 | -1.07 | 0.0411 | 4.51e-06 | F |
| 12 | L6/2417 | 1.29 | 1.40 | 0.18 | 3.91e-05 | P |
| 13 | L6/1335 | -0.14 | -0.26 | 0.0335 | 1.16e-05 | F |
| 14 | L1/2023 | 4.39 | 0.72 | 0.389 | 3.89e-06 | M |
| 15 | L1/2177 | 3.89 | -0.28 | 0.409 | 3.88e-06 | M |
| 16 | L1/1982 | 87.53 | 0.29 | 3.18 | 3.95e-06 | M |

**Figure 9:**
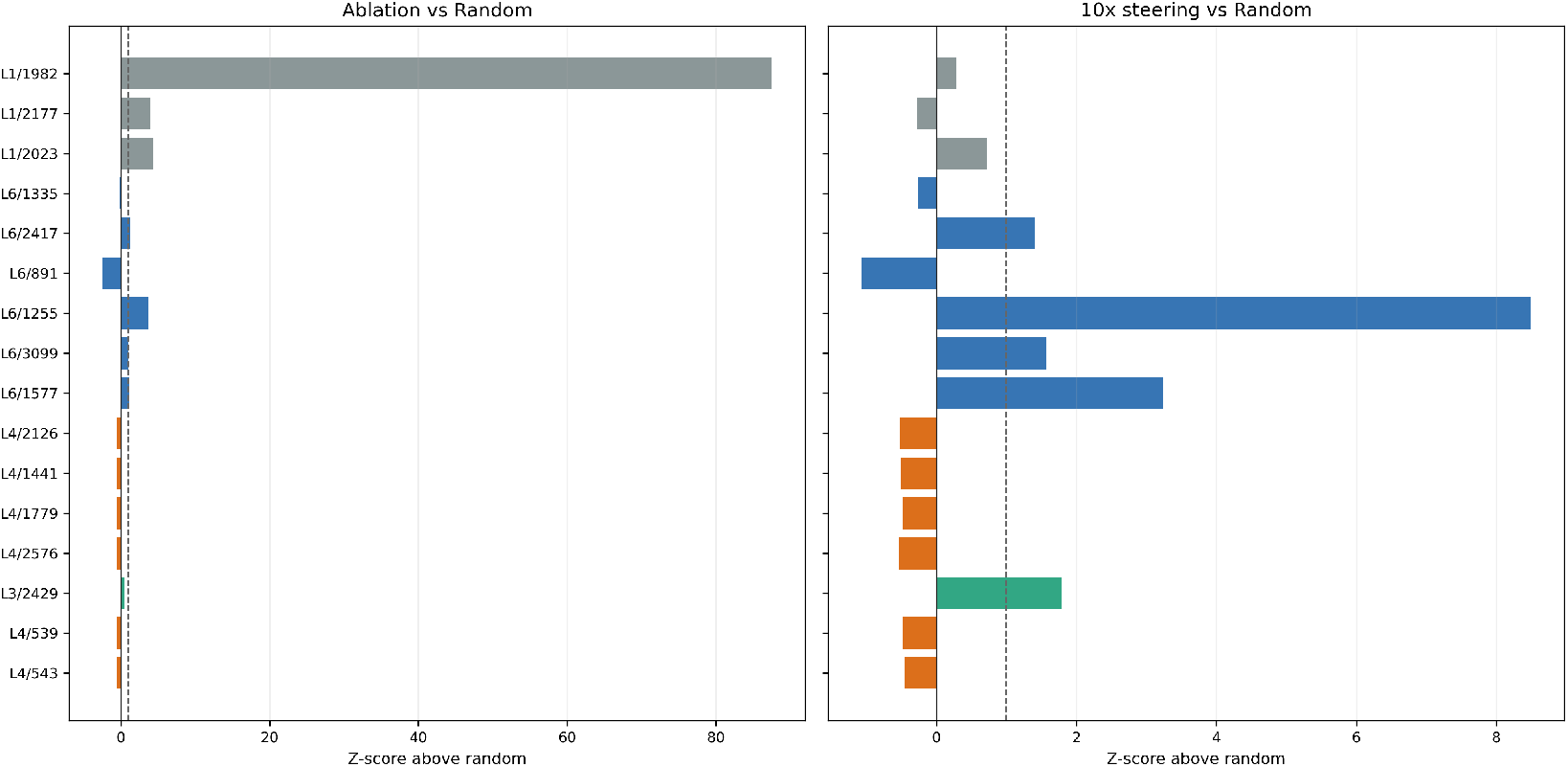
Probe interventions measured by standard deviations above random controls. The left figure is for ablation and the right figure is for 10× steering.

On the other hand, the nodes in layer 6 show the best results. L6/1255 produces the strongest probe effects in the whole circuit with +3.70 standard deviations for ablation and +8.50 standard deviations for 10× steering. Also, nodes such as L6/1577 and L6/3099 produced high positive values for ablation and steering. So, at least in this stage these layer 6 nodes pass, but it is important to note that L6/1255 with the strongest result fails in the cross-comparison stage.

### 4.6 Metagenome Validation

Table 8 and Figure 10 show the results for the metagenome dataset scan [23]. The metagenome results show similar patterns for the strongest and weakest nodes. The strongest nodes are zero or close to zero for the metagenome vs Class A ratio such as all layer 4 nodes being below 0.001 and the closest to 0. They are also zero for the activation threshold assessment where the nodes ranked 1 to 9 all return 0 meaning there are no metagenome sequences that rise above even the 75% activation threshold.

**Table 8:** Metagenome summary for all Class A circuit nodes.

| Rank | Node | Metagenome/Class-A ratio | >75th | >90th | >95th | Status |
| --- | --- | --- | --- | --- | --- | --- |
| 1 | L4/543 | 0.001 | 0 | 0 | 0 | P |
| 2 | L4/539 | 0.001 | 0 | 0 | 0 | P |
| 3 | L3/2429 | 0.000 | 0 | 0 | 0 | P |
| 4 | L4/2576 | 0.000 | 0 | 0 | 0 | P |
| 5 | L4/1779 | 0.000 | 0 | 0 | 0 | P |
| 6 | L4/1441 | 0.000 | 0 | 0 | 0 | P |
| 7 | L4/2126 | 0.000 | 0 | 0 | 0 | P |
| 8 | L6/1577 | 0.008 | 0 | 0 | 0 | P |
| 9 | L6/3099 | 0.211 | 0 | 0 | 0 | M |
| 10 | L6/1255 | 0.063 | 4 | 3 | 3 | M |
| 11 | L6/891 | 1.337 | 365 | 348 | 341 | F |
| 12 | L6/2417 | 1.026 | 185 | 143 | 136 | F |
| 13 | L6/1335 | 1.253 | 234 | 191 | 178 | F |
| 14 | L1/2023 | 0.989 | 183 | 178 | 148 | F |
| 15 | L1/2177 | 1.073 | 220 | 168 | 167 | F |
| 16 | L1/1982 | 0.984 | 41 | 30 | 29 | F |

**Table 9:** Framework Pass/Fail Table for All Class A Circuit Nodes.

| Rank | Node | Role | Benchmarking | Cross-Class | Family Breadth | Structural Grounding | Intervention | Metagenome |
| --- | --- | --- | --- | --- | --- | --- | --- | --- |
| 1 | L4/543 | strong candidate | P | P | P | P | F | P |
| 2 | L4/539 | strong candidate | P | P | P | M | F | P |
| 3 | L3/2429 | shared class feature | P | F | P | P | M | P |
| 4 | L4/2576 | strong candidate | P | P | P | P | F | P |
| 5 | L4/1779 | strong candidate | P | P | P | P | F | P |
| 6 | L4/1441 | strong candidate | P | P | P | M | F | P |
| 7 | L4/2126 | strongest discriminator, weak cross-class | P | F | P | M | F | P |
| 8 | L6/1577 | strong candidate | P | P | P | M | P | P |
| 9 | L6/3099 | shared class feature | P | F | P | M | P | M |
| 10 | L6/1255 | shared class feature | P | F | P | M | P | M |
| 11 | L6/891 | feature for leucine | M | F | F | F | F | F |
| 12 | L6/2417 | feature for alanine | M | F | F | F | P | F |
| 13 | L6/1335 | overall weak candidate | F | F | F | F | F | F |
| 14 | L1/2023 | feature for valine | F | F | F | F | M | F |
| 15 | L1/2177 | feature for glycine | F | F | F | F | M | F |
| 16 | L1/1982 | feature for glutamine | F | F | F | F | M | F |

P = Pass, M = Mixed, F = Fail.

**Figure 10:**
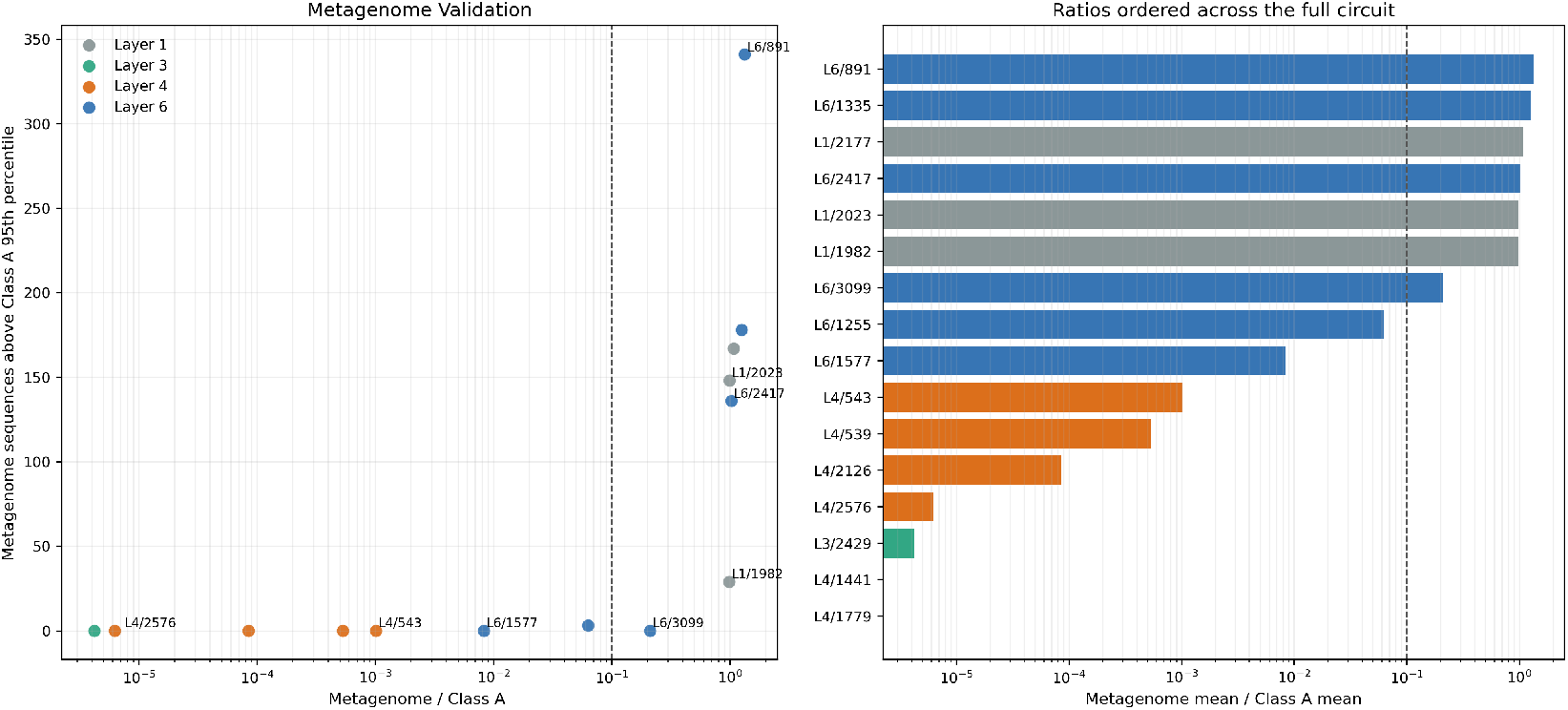
The left figure shows the metagenome vs Class A ratio versus the number of metagenome proteins going above the 95th percentile threshold. The right figure shows the same activation ratios ordered across the full circuit. In both figures, the vertical dotted line denotes zero. The strongest nodes cluster at the bottom-left corner of the plot with low metagenome vs Class A ratios and low amount of metagenome sequences that go above the threshold. On the other hand, the weakest are grouped at the opposite corner in the top-right corner.

However, the worst nodes have high metagenome vs Class A ratios and return many sequences for the activation thresholds. For example, the worst node in this stage is L6/891 with a metagenome vs Class A ratio of 1.337, 365 sequences over the 75% threshold, 348 sequences over the 90% threshold, 341 sequences over the 95% threshold. The ratio means that node L6/891 activates 133% more on metagenome sequences than Class A sequences and the threshold means that for each threshold there are more than 300 metagenome sequences that go above it. These results could mean that node L6/891 is not a feature for Class A *β*-lactamase but should be validated by the other stages.

## 5 Discussion

### 5.1 What the SAE and CLT Results Show Together

The InterPLM SAE (sparse autoencoder) analysis and ProtoMech CLT (cross-layer transcoder) analysis show similar, but slightly different results. The SAE results show that ESM-2 contains a strong separation between Class A and Class B at the final hidden layer. On the other hand, the CLT results also show that there is strong separation between Class A and Class B, C, and D, but that responsibility is spread between nodes across different layers with different roles.

### 5.2 Validation Framework Pass or Fail Results for Class A Circuit

Table 9 summarizes the results of the validation framework from Figure 1 as pass (P) or fail (F). There are two main outcomes from this table: no node passes every core stage and the combined framework catches different mistakes which usually were cross-class comparison failure or superficial patterns. For the first main outcome, even the best candidates that pass all other stages, L4/2576, L4/543, and L4/1779, fail the probe intervention stage. For the second main outcome, some examples include L3/2429 failing cross-class comparison, L6/2417 being a superficial pattern for alanine, and the layer 1 nodes being broad failures.

#### 5.2.1 Class A Circuit Roles and Layer-wise Analysis

Furthermore, through the validation framework, we can give three different roles to each node in the Class A circuit: a superficial feature for a specific amino acid, a feature shared among different *β*-lactamase classes, or a strong Class A feature candidate. These roles are shown in Table 9 as well. There also seems to be a pattern for the roles where layer 1 and other nodes ranked towards the bottom of Table 9 are superficial features for a specific amino acid, layer 6 has most of the shared class features, and layer 4 has most of the strong candidate nodes.

Together, these patterns in the Class A circuit roles support the theory found by previous research with SAEs and CLTs that ESM-2 has three distinct parts: the early layers find general features for residues, the middle layers are features for true structural features, and the last or late layers are features for specific residues [17]. So, this theory explains some of the results of the validation framework. For example, out of the six stages, the probe intervention stage gave the most interesting results because all the nodes with the role of strong candidate fail the probe intervention stage except for L6/1577. These results seem to support the notion that the later layers of a PLM find the most important details necessary for the final output[17].

### 5.3 Implications for AMR Surveillance

The current paper is not a usable surveillance system. The metagenome comparison shows why caution is necessary: superficial nodes are highly active on the metagenome dataset, and some of the strongest nodes for this stage fail in several other validation stages. However, if stronger validation becomes available, circuits discovered from CLTs could eventually support AMR surveillance [24]. So instead of using a single node, one could use a small validated circuit of nodes that have to agree across all validation stages.

### 5.4 Limitations

This paper has four main limitations:

1. **The paper evaluates a recovered circuit, not the full model exhaustively**. The main CLT analysis focuses on the 16 node Class A circuit to reduce computational load but looking at the full CLT might reveal even better Class A *β*-lactamase nodes that were not included in the circuit.
2. **The model is small**. ESM-2 8M is the smallest model in the ESM-2 family which makes the computational load easier especially for CLTs which scale exponentially based on the size of the base model [17]. However, larger ESM-2 models could reveal even more and better features for Class A *β*-lactamase within SAEs and CLTs.
3. **Structural grounding was not exhaustive**. There are more motifs for Class A *β*-lactamase than the three motifs that we looked at, so the structural grounding could have been stricter. Also, only the SxxK motif was found in all Class A while the other motifs have some variations within Class A although they are rare.
4. **Metagenome dataset was not the entire NMPFamsDB**. We only looked 5,000 of the over 70,000 sequences contained within the dataset from NMPFamsDB to reduce computational workload which this specific dataset itself is a subset of only bacteria where the full dataset is millions of sequences [23].

## 6 Conclusion

This paper showed that ESM-2 8M [22] contains strong candidate nodes or features that are able to separate Class A from Class B, C, and D *β*-lactamases [6] and that there are many strong candidate nodes or features rather than just one clear node within the Class A circuit. So, the main contribution in this paper is the validation framework where the six stages shown in Figure 1 help to distinguish strong candidate nodes from weak, superficial nodes. Together, these stages separate strong candidates from shared class features such as L3/2429 and L6/1255 and superficial patterns such as L6/2417 and the layer 1 nodes.

Several future directions are balancing the classes, analyzing the entire CLT instead of a recovered circuit, and training CLTs for larger ESM-2 models so that future work can replicate this work in larger ESM-2 models. There are already SAEs trained on the larger ESM-2 models [19]. Another future work could be to apply the validation framework to SAE features but CLTs seem to be the current standard for mechanistic interpretability [17]. For any practical AMR surveillance, these results would still need validation in the real world and the validation framework would need to be benchmarked against modern tools.

## A InterPLM and ProtoMech Scripts and Models

We used some scripts from InterPLM for the initial feature extraction and from ProtoMech for circuit discovery, benchmarking, and probe interventions: The scripts for InterPLM SAEs can be found here: https://github.com/ElanaPearl/InterPLM. The scripts for ProtoMech CLTs and circuits can be found here: https://github.com/amirgroup-codes/ProtoMech.

Also, the ESM-2 8M InterPLM SAE can be found here: https://huggingface.co/Elana/InterPLM-esm2-8m. The ESM-2 8M ProtoMech CLT can be found here: https://huggingface.co/ktalreja/ProtoMechModels/tree/main/CLT_L6_D3200

## B Supplementary Methods and Tables

### B.1 Metric Formulas

This subsection gives the mathematical definitions for the sequence-level metrics summarized in Chapter 3. Let *i* index proteins, *p* sequence positions, and *f* SAE features or CLT nodes. Let *y*_*i*_ ∈ {0, 1} denote the class label, let *I*_*c*_ denote the set of proteins in class *c*, and let *a*_*ipf*_ be the position-wise activation of feature or node *f* at position *p* in protein *i*.

#### B.1.1 Sequence-Level Mean Activation

For protein *i* of length *L*_*i*_, the per-sequence mean activation of node or feature *f* is

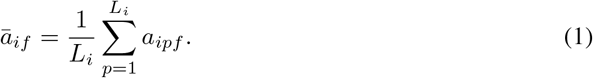

#### B.1.2 Cohen’s *d*

For two comparison classes *c*_1_ and *c*_2_, separation is summarized with

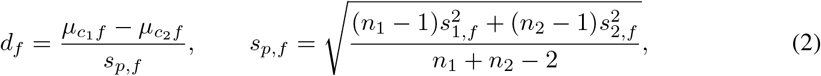

where *µ, s*^2^, and *n* denote the class mean, variance, and sample size for feature *f* .

#### B.1.3 Area Under the Receiver Operating Characteristic Curve

Let 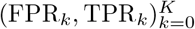 denote the points on the empirical ROC curve ordered by increasing false-positive rate. AUROC is computed:

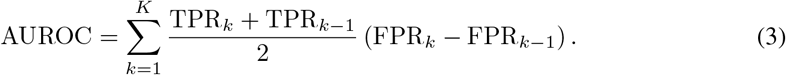

#### B.1.4 Spearman’s *ρ*

For paired observations 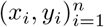 with rank variables *R*_*i*_ = rank(*x*_*i*_) and *S*_*i*_ = rank(*y*_*i*_), Spear-man’s rank correlation is:

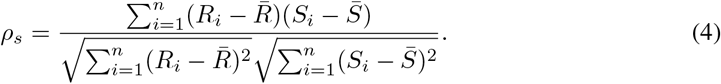

The reported *p*-values test the null hypothesis *ρ*_*s*_ = 0.

#### B.1.5 Intervention Shifts

If *ℓ*_*i*_ and *p*_*i*_ denote the baseline probe logit and probability for sequence *i*, and 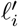 and 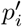 the post-intervention values, we report:

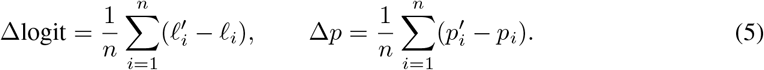

#### B.1.6 Metagenome Ratios and Overlap

Let ℳ be the metagenome set, the verified Class A set, and *T*_*N*_ (*f*) the set of top-*N* proteins ranked by node *f* . The mean-activation ratio and top-*N* overlap are:

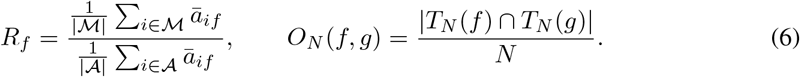

#### B.1.7 Metric Precedent in InterPLM and ProtoMech

Table 10 summarizes the core standard metrics used in this paper and their precedent in InterPLM and/or ProtoMech.

**Table 10:** Core metric precedent in InterPLM and ProtoMech.

| Metric | Used here for | Precedent |
| --- | --- | --- |
| AUROC | SAE discrimination, CLT benchmarking, family breadth, and cross-class transfer | InterPLM [19]; original AUROC reference: Hanley and McNeil [25] |
| Cohen’s $d$ | SAE and CLT effect-size summaries | InterPLM [19] |
| Welch’s $t$-test | SAE mean-difference testing | InterPLM [19]; original reference: Welch [26] |
| Spearman’s $\rho$ | Active site distance | InterPLM [19] and ProtoMech [17] |

**Table 11:** Full Class A circuit benchmark summary, ordered by layer.

| Rank | Node | A vs B | A vs C | A vs D | Cohen's $d$ | Min fam | Global rank | Status |
| --- | --- | --- | --- | --- | --- | --- | --- | --- |
| 16 | L1/1982 | 0.234 | 0.093 | 0.219 | -1.03 | 0.003 | 18716 | F |
| 14 | L1/2023 | 0.304 | 0.531 | 0.338 | -1.27 | 0.186 | 18044 | F |
| 15 | L1/2177 | 0.255 | 0.326 | 0.597 | -0.83 | 0.169 | 18325 | F |
| 3 | L3/2429 | 0.998 | 0.206 | 0.963 | 19.01 | 0.996 | 11 | P |
| 2 | L4/539 | 0.999 | 0.993 | 0.945 | 18.87 | 1.000 | 2 | P |
| 1 | L4/543 | 1.000 | 0.980 | 0.915 | 19.04 | 1.000 | 1 | P |
| 6 | L4/1441 | 1.000 | 0.999 | 0.946 | 12.16 | 1.000 | 30 | M |
| 5 | L4/1779 | 1.000 | 0.999 | 0.916 | 14.28 | 1.000 | 17 | P |
| 7 | L4/2126 | 1.000 | 0.600 | 0.793 | 13.27 | 1.000 | 38 | M |
| 4 | L4/2576 | 1.000 | 0.967 | 0.970 | 15.84 | 1.000 | 12 | P |
| 11 | L6/891 | 0.781 | 0.848 | 0.758 | 1.01 | 0.143 | 785 | F |
| 10 | L6/1255 | 0.997 | 0.000 | 0.815 | 4.38 | 0.999 | 332 | M |
| 13 | L6/1335 | 0.744 | 0.151 | 0.214 | 0.91 | 0.635 | 1303 | F |
| 8 | L6/1577 | 0.999 | 0.999 | 0.993 | 4.79 | 0.999 | 103 | P |
| 12 | L6/2417 | 0.771 | 0.723 | 0.871 | 1.24 | 0.696 | 795 | M |
| 9 | L6/3099 | 0.999 | 0.999 | 0.738 | 4.60 | 1.000 | 153 | M |

**Table 12:** Per-family AUROC for all Class A circuit nodes against the full Class B pool.

| Rank | Node | TEM | SHV | CTX-M | KPC | GES | Status |
| --- | --- | --- | --- | --- | --- | --- | --- |
| 1 | L4/543 | 1.000 | 1.000 | 1.000 | 1.000 | 1.000 | P |
| 2 | L4/539 | 1.000 | 1.000 | 1.000 | 1.000 | 1.000 | P |
| 3 | L3/2429 | 1.000 | 1.000 | 0.996 | 1.000 | 1.000 | P |
| 4 | L4/2576 | 1.000 | 1.000 | 1.000 | 1.000 | 1.000 | P |
| 5 | L4/1779 | 1.000 | 1.000 | 1.000 | 1.000 | 1.000 | P |
| 6 | L4/1441 | 1.000 | 1.000 | 1.000 | 1.000 | 1.000 | P |
| 7 | L4/2126 | 1.000 | 1.000 | 1.000 | 1.000 | 1.000 | P |
| 8 | L6/1577 | 1.000 | 0.999 | 1.000 | 1.000 | 1.000 | P |
| 9 | L6/3099 | 1.000 | 1.000 | 1.000 | 1.000 | 1.000 | P |
| 10 | L6/1255 | 1.000 | 0.999 | 1.000 | 1.000 | 1.000 | P |
| 11 | L6/891 | 0.813 | 0.967 | 0.753 | 0.990 | 0.143 | F |
| 12 | L6/2417 | 0.696 | 0.787 | 0.792 | 0.992 | 0.789 | F |
| 13 | L6/1335 | 0.636 | 0.948 | 0.861 | 0.635 | 0.640 | F |
| 14 | L1/2023 | 0.186 | 0.273 | 0.337 | 0.410 | 0.386 | F |
| 15 | L1/2177 | 0.178 | 0.169 | 0.170 | 0.600 | 0.311 | F |
| 16 | L1/1982 | 0.046 | 0.027 | 0.003 | 0.790 | 0.042 | F |

As for the other metrics in this paper, they are usually derived from these core metrics.

### B.2 Class A Circuit Tables

